# Facial expression recognition is modulated by approach–avoidance behavior

**DOI:** 10.1101/2024.05.21.594616

**Authors:** Yugo Kobayashi, Hideki Tamura, Shigeki Nakauchi, Tetsuto Minami

**Affiliations:** Department of Computer Science and Engineering, Toyohashi University of Technology, Toyohashi, Aichi, Japan

**Keywords:** Facial expression, approach–avoidance behavior, happy, angry, fearful, facial expression recognition

## Abstract

Facial expression recognition influences approach–avoidance behaviors, but do approach–avoidance behaviors affect facial expression recognition? We conducted psychophysical experiments to this end, indicating a reverse causal relationship. In a virtual reality space, 3D face stimulus facial expressions varied on seven levels—from happy to angry in Experiments 1 and 3 and from happy to fearful in Experiment 2. Participants were asked to perform according to one of the following conditions in response to the stimulus. Participants 1) approached (one-meter forward), 2) avoided (one-meter backward), 3) were approached by, or 4) were avoided by the 3D model. Then, participants selected facial expressions. Experiment 1 revealed that participants recognized the face as angrier when they avoided it rather than when it avoided them. Experiment 2 showed that participants recognized the face as happy when approaching and fearful when avoiding, irrespective of who acted. Experiment 3 revealed that participants recognized the face as angrier when the face approached them rather than when they approached if both parties were physically close. Accordingly, approach–avoidance behavior changes facial expression recognition, indicating a reverse causal relationship. We posit that unconscious learning rooted in biological instincts creates this connection.

## Introduction

Communication is an essential aspect of modern society. Human-to-human exchanges—including facial expressions and body language—involve both verbal and nonverbal cues. Cognitive science and psychology have made significant advances in the study of facial expressions. Facial expressions and behaviors influence each other. Specifically, emotional recognition influences stepping forward and backward; that is, by an approach–avoidance behavior. For example, moving toward a smiling person is faster than toward an angry person [1]. Additionally, individuals tend to maintain a greater distance from an angry person than from a smiling one [2,3].

Approach–avoidance behavior toward facial expressions has been investigated from various aspects such as “individual characteristics,” “physiological indices,” and “motivation” using various methods such as “actual walking” and “pseudo-approach–avoidance with lever.” Humans tend to approach positive facial expressions and avoid negative facial expressions [4], which is modulated by individual characteristics and sex. For example, when individuals exhibit depressive symptoms, the motivation of approach–avoidance behavior for facial expressions becomes weaker [5]. Additionally, interpersonal-sensitive individuals approach faces with happy and neutral expressions faster than avoidance compared to other individuals [6]. Moreover, skin potentials have shown that humans feel threatened by faces nearby approaching with angry facial expressions [7]. Other studies have suggested that humans automatically trigger a defensive response to stimuli approaching with angry facial expressions, bypassing other cognitive processes [8]. Further, a previous study that used arm bending exercises and a smartphone to assess approach–avoidance behavior reconfirmed that people are more likely to approach smiling faces and avoid angry faces [9].

However, no studies have examined how physical state or behavior influence facial expression recognition; that is, whether a reverse causal relationship exists between approach– avoidance behaviors and emotion recognition. This study answers the following question: “Do approach–avoidance behaviors affect facial expression recognition?” Previous studies have the confirmed original relationship [1–3,10,11]. Some are based on biological instincts [12,13]. Therefore, if there is such a strong relationship between behavior and cognition, it is not surprising that a reverse causal relationship is also established.

First, pleasant or desirable situations—such as erotic stimuli, nature, families, and foods— elicit approach behaviors. The approach is motivated by the appetitive system, which is facilitated by positive experiences. The presence of the appetitive system is physiologically supported by a modest initial deceleration and mid-interval acceleration of the heart rate (skin conductance increases at high excitability) [12]. Why humans are more likely to approach a smiling person than an angry one is easy to understand [1,2,10,11]. Similarly, humans approach pleasant images quicker than unpleasant ones [14]. Thus, approaching strongly connects positive things at the fundamental level with biological instincts.

Second, dangerous or threatening situations such as attacks on humans or animals, pictures of mutilated bodies, and contamination elicit avoidance behaviors. The avoidance is motivated by the defense system, which is facilitated by negative things. The defense system is physiologically supported by skin conductance increases, sustained heart rate deceleration (but heart rate acceleration for imminent threats), and so on (depending on context) [12]. Reportedly, along with using body movements, operating stimuli on displays, or manipulating shapes resembling oneself through levers can also promote avoidance tendencies toward anger or unpleasant images [15,16]. Avoidance connects to negative things similarly to the relationship between approaching positive things.

Third, the literature regarding social embodiment informs this research question. Our body movements and positions provide cues regarding our subjective state. For example, arm flexion and extension can be understood as an approach–avoidance behavior and self-generated bodily cues that reveal the detection time for positive and negative words, respectively [17,18]. In a previous study, participants swiped different positive and negative pictures closer or away from themselves [19]. These mechanisms have not yet been clarified, although the effects of changes in neuroendocrine levels and mirror neurons have been discussed; however, there are many reports of cognition changing with physical condition [20]. Those direct actions support why we connect approach–avoidance behavior with emotional experiences. Therefore, we posit that approach–avoidance behaviors can provide signals for facial expression recognition.

We conducted three experiments using approach–avoidance behaviors, which are expected to exhibit a strong motivating effect of revealing the impact of one’s actions on facial expression recognition. **Figure 1** presents a brief overview of this study. Specifically, participants with a head-mounted display were instructed to perform according to one of the following four conditions in a virtual reality (VR) space. They 1) approached (one-meter forward), 2) avoided (one-meter backward), 3) were approached by, or 4) were avoided by a 3D face model. Thereafter, they were asked to discriminate the facial expressions from two choices: happy or angry and happy or fearful in Experiments 1 and 2, respectively. Additionally, Experiment 3 decreases the distance between participants and the 3D face model to very whether this relationship changes with distance. The angry and happy pairs in Experiment 1 were determined based on previous studies [1,2]. In Experiment 2, we used happy and fearful expressions. Fear is the opposite of anger. If there is a difference in facial expression recognition between the condition in which participants move and the condition in which the 3D model moves, we can conclude that it is not owing to changes in distance, but owing to participants’ own movement (or the movement of others).

**Figure 1.**
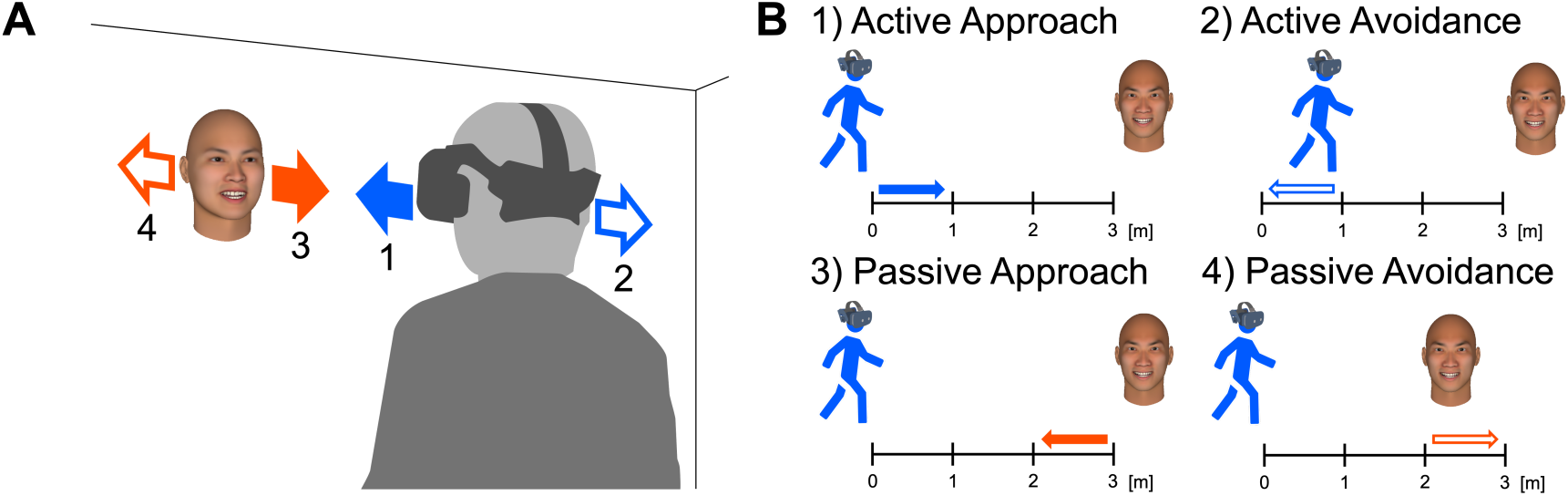
Experiment overviews. **A)** Sample image of a participant and face model in virtual reality in the experiment. Notably, the face model is located in the virtual reality space with a gray background (not augmented reality with real-world background). The person on the right is one of the authors, not a participant, and has given us permission to use this image as an example. **B)** Positional relationships between participants and the face model in each condition.

In Experiment 1, we hypothesized that it would be more likely to perceive a 3D model’s facial expressions as smiling when either participants or the 3D model face approached each other and as anger when participants avoided the 3D model. This hypothesis is based on the fact that participants are more likely to approach an opponent with a smiling face and to avoid an opponent with an angry face [1–3,10,11]. In Experiment 2, we hypothesized that we would be more likely to perceive a 3D model’s facial expressions as fearful when the model avoided participants. This hypothesis is based on our idea that we expect that others who move away from us appear to feel threatened by us.

## Experiment 1

We first examined whether approach–avoidance behavior affects the recognition of happy and angry facial expressions. Previous studies [12–14] have suggested that participants approach benefits and avoid harm. Therefore, we hypothesized that people would easily recognize a friendly “happy” expression and approach it and a threatening “angry” expression and avoid it. Using the VR with a head-mounted display (HMD) to present the stimuli releases the body constraints associated with using a monitor. Our methodology is advantageous compared to traditional research methods in the cognitive science field that use monitors exclusively.

## Methods

### Participants

Twenty-four men participated in the experiment (mean age = 23.25 years, *SD* = 0.83). Two participants were excluded from the subsequent analysis owing to being outlier points of subjective equality (PSE) value wherein the data deviated by two or more standard deviations from the mean in any experimental condition. The sample size was determined by performing a statistical power analysis using G*Power (version 3.1.9.7) [21,22] for a within-participants two-factor analysis of variance interaction with the direction condition (approach or avoidance) and the agency condition (active or passive). The parameters are as follows: effect size of f = 0.25, α = .05, and power = 0.80. Written informed consent was obtained from all participants after explaining to them the procedural details. The Committee for Human Research of the Toyohashi University of Technology approved the experimental procedures, which were conducted in accordance with the Declaration of Helsinki. We did not pre-register the experiment.

### Apparatus

The experimental stimuli were presented using an HMD, the HTC VIVE Pro Eye (resolution = 1440×1600 pixels per eye, refresh rate = 90 Hz, field of view = 110°), along with a Windows 10 PC (equipped with a Core i7-11700 processor and 32 GB RAM). The software used for the experiment were Unity 2021.1.25f1 and Steam VR 1.24.6. A numeric keypad was used to record participant responses. To obtain coordinate information in the VR space, one VIVE Tracker (3.0) was attached to the participant’s waist, along with four base stations (Steam VR Base Station 2.0) at the corners of the experimental area.

### Stimuli

Using the FaceGen Modeller Core 3.29 software, which generates a 3D model of the human face, we created 56 types of stimuli comprising facial expressions at seven levels. We then modified the parameters of “Expression Anger” from two to eight and “Expression SmileOpen” from eight to two to generate 3D models of the seven facial expression levels (**Figure 2A**). We refer to the seven expression levels as “Angry Levels 2–8.” The facial expressions generated by the FaceGen were recognized by participants as intended [23,24]. We then randomly generated eight individual faces with the “East Asian racial group” and “Any sex” settings. Therefore, the total number of stimuli was 56 (seven levels of facial expression × eight individual faces). Notably, the Identity 7 face was the same as the Identity 6 one during the experiment owing to a software malfunction.

**Figure 2.**
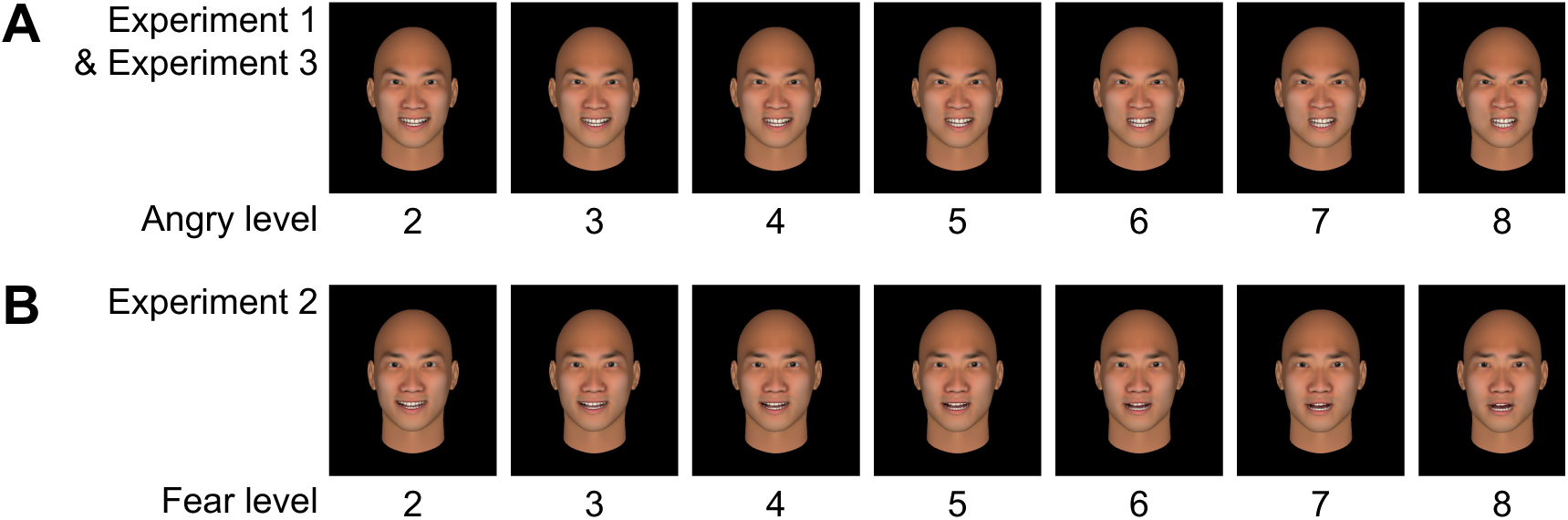
The model’s seven facial expression levels were modified between two expressions: **A)** happy and angry in Experiments 1 and 3. **B)** happy and fearful in Experiment 2.

The vertical and horizontal face stimuli sizes were 28.6 and 19.1 cm on average, respectively. At a distance of two meters, the average horizontal and vertical visual angles were 8.2° and 5.5°, respectively. At a distance of three meters, the average horizontal and vertical visual angles were 5.5° and 3.7°, respectively. These measurements were manually performed using Blender’s measuring tool. The distance was measured in Unity units, with one unit defined and converted as equal to one meter.

### Procedure

The procedure is illustrated in **Figure 3A**. Participants wore an HMD and a tracker on their waist, held a keypad in their hands (**Figure 3B**), and stood in a VR with a gray background (25 cd/m^2^). The sequence of participants’ actions in one trial was as follows. 1) The participant moved to the center of the room using a white square on the ground as a reference point. 2) The fixation point was presented for one second. 3) Text instruction was presented for one second. 4) The facial stimuli were presented until the participant completed the action. 5) The response screen was displayed.

**Figure 3.**
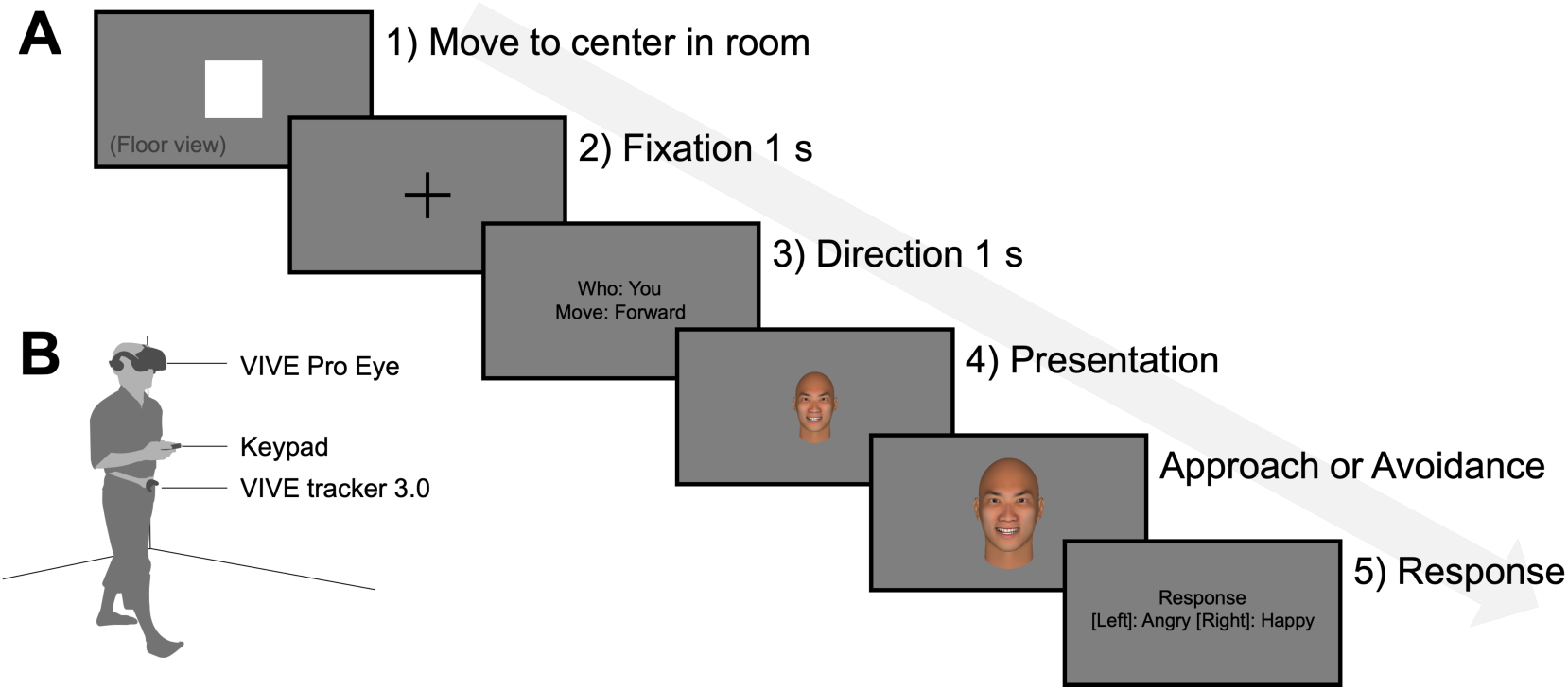
The procedure of Experiment 1. **A)** Paradigm. **B)** Participant’s view during the task. Notably, the person is one of the authors, not a participant, and has given us permission to use the image as an example.

The task had four conditions: “approach or avoidance” × “participant or facial stimuli moving.” If the text instruction was “Who: Avatar,” the facial stimuli were enacted by the model; if it was “Who: You,” the participant had to act. The actions are indicated as “Move: Forward” for approach and “Move: Backward” for avoidance. Facial stimuli were presented three meters away from the participant in the approach condition and two meters away in the avoidance condition, followed by a one-meter movement. The distance between the participant and the face model fit within the range of a social distance of 1.2–3.6 m [25]. Notably, the initial distance differed between the approach and avoidance conditions; however, the total visual information in the retina was identical for both conditions. Whether the participant or the face model is approaching, the movement is from three meters to two meters and the size of stimulus transferred on the retina is the same. For avoiding, the range is between two and three meters regardless of whether the participant or the model moves.

Stimuli were presented at approximately the same position as the height of the participant’s face by referring to the HMD coordinates. The facial stimuli’s movement speed was 0.8 m/s. Participants responded whether the face stimulus’ facial expression was “Happy” or “Angry” using a numeric keypad. Four sessions—with 56 trials per session—were conducted. All stimuli and conditions were executed in random order. Before the experiment, participants completed a practice task comprising eight trials (two trials per each of the four conditions) using the same conditions and maximum/minimum levels of angry stimuli as in the main experiment.

### Data analysis

Data were analyzed using MATLAB R2021b. To eliminate the effects of unexpected artifacts that occurred during the experiment, trials were excluded using the following criteria. 1) The tracker’s coordinates could not be obtained for the condition that participants acted. 2) The participants moved more than 50 cm to the left or right. 3) The participants moved more than 50 cm in the opposite direction of the instruction. 4) Either the response or action took more than 10 s. 5) The participant did not see the front within a 30° visual angle for 90% of the stimulus presentation time. Thus, on average, 1.05 trials per participant were excluded (standard error 0.43).

The proportion of trials wherein participants responded with “angry” was determined for each experimental participant, condition, and anger level. The psychometric functions representing the angry proportion were fitted using a logistic function and the Palamedes Toolbox [26]. Subsequently, the PSE was determined for each participant and condition. High PSE values indicated that participants were more likely to respond to the facial model’s expression as “happy,” while low values indicate that they were more likely to respond to it as “angry.” Statistical analyses were performed using JASP (Version 0.16.3) [27]; further, Bonferroni correction was used for multiple comparisons.

## Results

**Figure 4A** presents the PSE values for each condition. The PSE values were as follows: active approach (mean = 6.144, SE = 0.243), passive approach (mean = 6.197, SE = 0.276), active avoidance (mean = 5.498, SE = 0.256), and passive avoidance (mean = 6.452, SE = 0.431). The all condition mean was 6.073. We performed a two-way repeated-measures analysis of variance for the direction condition (approach/avoidance) and agency condition (active/passive). The direction condition exhibited no main effect (*F*(1,21) = 0.528, *p* = .475, η_p_^2^ = 0.025). By contrast, the agency condition exhibited a significant main effect (*F*(1,21) = 5.299, *p* = .032, η_p_^2^ = 0.201). This means that anger was more easily recognized when participants moved than when the face model moved.

**Figure 4.**
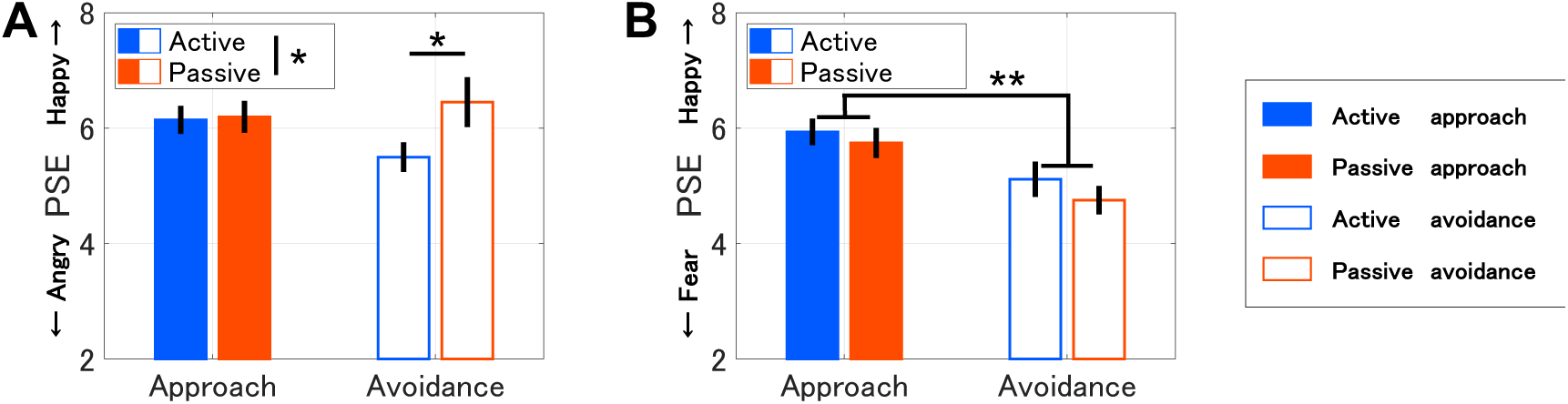
The results of Experiments 1 and 2. “Active” represents the condition wherein participants acted, and “passive” represents a condition wherein the facial stimulus acted. PSE: point of subjective equality. **A)** Experiment 1: The vertical axis indicates a PSE with smaller values representing easier anger recognition. **B)** Experiment 2: The vertical axis indicates a PSE with smaller values representing easier fear recognition. The error bars represent the standard error and asterisks indicate significant differences: \**p* < .05, \*\**p* < .01.

Moreover, there was a significant interaction (*F*(1,21) = 4.457, *p* = .047, η_p_^2^ = 0.175). The post-hoc tests revealed a difference between active avoidance and passive avoidance (*t*(21) = 3.122, *p* = .019, Cohen’s *d* = 0.612). This means that anger was more easily recognized when participants avoided the stimuli than when the stimuli avoided them.

## Discussion

In Experiment 1, participants responded to the facial expressions of a 3D facial model after the participant or the model performed approach–avoidance behavior in VR space using HMD and Tracker. The model’s facial expressions were changed in seven levels, from happy to angry. Previous studies wherein facial expression recognition affected approach–avoidance behavior demonstrated that compared to seeing a happy facial expression, seeing an angry expression results in the individual maintaining a larger interpersonal distance from another [2,3] and a slower approach [1]. Further, other studies have suggested a fast avoidance response to threats [12–14,16,28]. These studies support the association between anger and avoidance, indicating that the purpose is maintaining distance from the threatening other. Therefore, this difference suggests that anger is not recognized more easily under conditions wherein one individual can maintain the distance from another, and the purpose can be fulfilled without moving oneself.

The results of Experiment 1 allow us to consider the hypothesis of Experiment 2 that the facial expressions of the others who move away from us are recognized as more likely fearful. Experiment 1 indicated that when participants avoid the other, the other’s facial expression is recognized as more likely angry (threatening facial expression). If participants felt threatened and avoided the other, we can imagine that they would have fearful facial expressions (threatened). If we consider this situation from the standpoint of the other, participants who move away from us will have fearful facial expressions. Thus, In Experiment 2, we attempted to gain further insight by using fear facial expressions, which are the same negative and opposite position of anger in Experiment 1.

## Experiment 2

In Experiment 2, we hypothesized that a person avoiding others would be recognized as having a fearful facial expression.

## Methods

### Participants

We determined the sample size in the same way as in Experiment 1, and 24 men participated (mean age = 22.00 years, *SD* = 1.22). Four participants were excluded from the subsequent analyses based on the same criteria as in Experiment 1. Written informed consent was obtained from all participants after explaining the procedural details to them. The Committee for Human Research of the Toyohashi University of Technology approved the experimental procedures, which were conducted in accordance with the Declaration of Helsinki. We did not pre-register the experiment.

### Apparatus

The apparatus was the same as that used in Experiment 1.

### Stimuli

**Figure 2B** presents the stimuli used in Experiment 2, which were the same as in Experiment 1, except that the angry expressions were changed to fearful ones.

### Procedure

The procedures were identical to those used in Experiment 1, except that the choices were happy and fearful instead of happy and angry.

### Data analysis

We performed the same analysis as in Experiment 1. The average number of excluded trials was 0.95 per participant (standard error 0.22). Each participant’s PSE was calculated based on the percentage of participants who answered “fear” in response to the stimulus’ fear level for each condition.

## Results

**Figure 4B** presents the PSE values for each condition. The PSE values were as follows: active approach (mean = 5.936, SE = 0.231), passive approach (mean = 5.744, SE = 0.262), active avoidance (mean = 5.116, SE = 0.306), and passive avoidance (mean = 4.753, SE = 0.247). The all condition mean was 5.387. A main effect was observed for the direction condition (*F*(1, 19) = 12.767, *p* = .002, η_p_^2^ = 0.402), meaning that avoidance and approach behaviors made the face model easier to recognize as fearful and happy, respectively.

By contrast, no main effect was observed for the agency condition (*F*(1, 19) = 1.564, *p* = .226, η_p_^2^ = 0.076), and no interaction was observed (*F*(1, 19) = 0.176, *p* = .680, η_p_^2^ = 0.009).

## Discussion

In Experiment 2, as in Experiment 1, participants responded to the facial expressions of a 3D face model after performing approach–avoidance behavior. The model’s facial expressions were changed in seven levels, from happy to fearful.

The results suggest that we were more likely to recognize the other’s facial expression as happy when the distance between us and the other was small, and we were more likely to recognize the other’s facial expression as fearful when the distance between us and the other was large. These results are consistent with the trend reported by previous studies in that comparing a smiling person to a fearful one makes the former more approachable [4]; that is, participants approached positive expressions and avoided negative ones.

However, no difference between agency condition in avoidance was observed. One reason is that fear is not invariably associated with avoidance. Fearful expressions are approached more quickly than angry ones, although both are negative [15,29,30]. Additionally, the explanation for the faster approach toward fear is twofold: sympathy toward frightened individuals [15] and a wide-eyed look resembling a child and attracting attention [30]. However, fear is also approached more quickly than positive expressions, such happiness. This faster avoidance occurs possibly because a fearful expression conveys the presence of a threat in the surrounding environment, from which observers may flee [31]. Thus, the aforementioned reports help explain why the passive avoidance condition did not enhance the fearful expression in this study.

Whether a person associates fear with approach or avoidance depends on whether the other individual is part of their ethnic group [31], whether the observer of the expression is sociable or individualistic [29], and the specific situation [32]. Therefore, examining whether fear is associated with approach or avoidance compared to other negative expressions is necessary.

## Experiment 3

In Experiment 3, we changed the distance from stimuli to participants to consider the effect of distance, in which the range is between 0.2 and 1.2 m. Because it is possible that the results of Experiment 1 were because the stimuli were more easily perceived as overall positive expressions. We hypothesized that anger was more easily recognized when the stimulus approached participants than when participants approached the stimulus and when participants avoided the stimulus than when the stimulus avoided participants. A previous study suggests that individuals feel uncomfortable when approaching angry others up to approximately 1.2 m and when being approached by angry others up to 1.6 m [2]. We expected a difference between participants approaching and the stimulus approaching, which was not seen in Experiment 1, as the range was outside 1.6 m (the range between 2.0 and 3.0 m).

## Methods

### Participants

We determined the sample size in the same way as in Experiment 1, and 24 men participated (mean age = 22.46 years, *SD* = 1.26). Four participants were excluded from the subsequent analyses based on the same criteria as in Experiment 1. Written informed consent was obtained from all participants after explaining the procedural details to them. The Committee for Human Research of the Toyohashi University of Technology approved the experimental procedures, which were conducted in accordance with the Declaration of Helsinki. We did not pre-register the experiment.

### Apparatus

The apparatus was the same as that used in Experiment 1.

### Stimuli

**Figure 5** presents the stimuli used in Experiment 3, which were the same as in Experiment 1.

**Figure 5.**
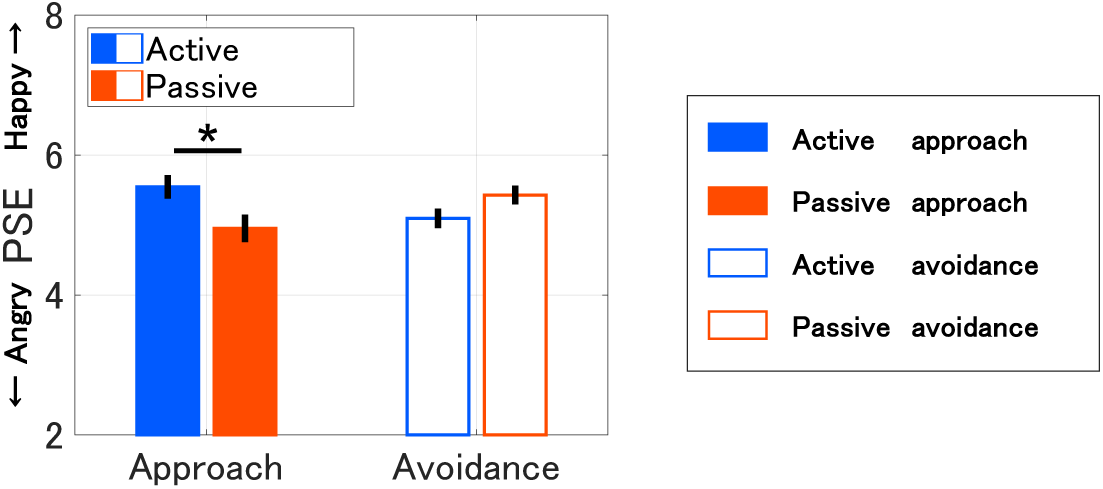
The results of Experiment 3. The format is the same as in Figure 4A.

### Procedure

The procedures were identical to those used in Experiment 1, except that the distance between the participant and the face model changed from 2.0–3.0 m to 0.2–1.2 m. This distance was determined as when participants felt uncomfortable [2] with the other under both conditions when the other approached and when participants approached. Additionally, this range is close to the experiment of Candini and colleagues [33].

### Data analysis

We performed the same analysis as in Experiment 1. The average number of excluded trials was 0.42 per participant (standard error 1.15).

## Results

**Figure 5** presents the PSE values for each condition. The PSE values were as follows: active approach (mean = 5.545, SE = 0.168), passive approach (mean = 4.951, SE = 0.195), active avoidance (mean = 5.096, SE = 0.139), and passive avoidance (mean = 5.428, SE = 0.135). The all condition mean was 5.255. The mean for all conditions in Experiment 3 was lower than in Experiment 1 (6.073).

Although no main effect was observed for the agency condition (*F*(1, 19) = 0.855, *p* = .367, η_p_^2^ = 0.043) and the direction condition (*F*(1, 19) = 0.007, *p* = .935, η_p_^2^ = 3.622 × 10^-4^), interaction was observed (*F*(1, 19) = 12.855, *p* = .002, η_p_^2^ = 0.404). The post-hoc tests revealed a difference between the self-approach and the other-approach (*t*(19) = 3.100, *p* = .022, Cohen’s *d* = 0.733). This means that anger was more easily recognized when the stimuli approached participants than when participants approached them.

However, unlike in Experiment 1, no significant differences between active avoidance and passive avoidance were observed (*t*(19) = 1.734, *p* = .547, Cohen’s *d* = 0.410). This means that anger was more easily recognized when participants avoided the stimuli than when the stimuli avoided participants but only in the situation in which the other was far from participants.

## Discussion

In Experiment 3, as in Experiment 1, participants responded to the facial expressions of the face 3D model after the approach-avoidance behavior. The distance between the model and the participant was smaller than in Experiment 1.

In Experiment 1, in which the distance was greater, the stimulus expression was more likely to be recognized as happy than in Experiment 3. Thus, it is possible that the reason the PSEs were larger and facial expressions were more easily perceived as positive in Experiment 1 was owing to the greater distance.

Results suggest that anger was more easily recognized when the stimuli approached participants than when participants approached them. We consider this because participants felt threatened by the other approaching. Previous studies—wherein facial expression recognition affected approach behavior—have demonstrated that stimuli approaching participants resulted in one individual maintaining a larger interpersonal distance from the other than when participants approached the stimuli [2,34,35]. In addition, other studies have stated that participants felt threatened by physiological indicators and defense responses [7,8]. These studies consider the cause to be that participants could not control the distance from the other [34]. Results in Experiment 1 suggest that anger was more easily recognized when participants avoided the stimuli than when the stimuli avoided participants but results in Experiment 3 suggest only in the situation in which the other was far from participants. We consider the effect of avoidance to influence situations to be visually ambiguous and influenced by predictions.

## General discussion

We conducted three experiments to determine whether approach–avoidance behavior affects facial expression recognition. In Experiment 1, we hypothesized that people would easily recognize a friendly “happy” expression when approaching it and a threatening “angry” expression when avoiding it. In Experiment 2, we hypothesized that a person’s avoidance behavior would be recognized as fear. Consequently, we can easily recognize that another is fearful when we or the other exhibits avoidance. In Experiment 3, we hypothesized that people would easily recognize a threatening “angry” expression when the other approached them. Experiments 1 and 2 revealed that participants recognized the face model’s expression as positive when approaching it and as negative when avoiding it. However, when the distance is small, it is easier to recognize the other as angry when they approach. Therefore, depending on distance, the behavior that influences facial expression recognition changes. This indicates the cognitive processing of not only the conventional relationship of “facial expression recognition affecting approach–avoidance behavior” but also the reverse causal relationship of “approach–avoidance behavior affecting facial expression recognition.”

A similar phenomenon is the “space–valence metaphor” [36,37,38], a cognitive characteristic that associates “up” and “handed side” with desirable things and “down” and “non-dominant side” with undesirable things. The fact that people perceive the dominant hand side as better suggests that spatially affective metaphors are the result of acquired learning [36]. Approach and avoidance are strongly associated with positive and negative stimuli, respectively, with respect to physiological aspects [12–14] and are unconsciously affected by others’ facial expressions [4]. Therefore, humans would have unconsciously internalized the connection between approach– avoidance and facial expressions. Therefore, we assume that unconsciously learning to approach or avoid an individual based on their expressions has led to approach–avoidance behavior’s influence on facial expression recognition.

We investigated only the relationship between positive and negative expressions. Remarkably, social factors, including sympathy and challenges, also influence approach–avoidance behavior [4,15,29,30]. Regarding sympathy, sad and fearful expressions make the approach behavior faster than other negative expressions such as anger or disgust [4,15]. As such, we need to investigate which social factors affect the approach–avoidance behavior by directly comparing negative expressions (e.g., anger and fear).

This study has several limitations. First, stimuli were presented using a 3D model of a face with no hair or body. Adding a body to the model can provide valuable insights because approach– avoidance behavior is influenced by a person’s physical state [30] and social factors [39]. Thus, to obtain more robust results, stimulus quality can be improved by adding hair and a body to the 3D models, as well as by providing a more realistic environment.

Second, a non-motion condition would help clarify the relationship between actions and facial expression recognition. We interpreted the results of Experiment 2 as the approach behavior being more easily recognized as “happy” than the avoidance behavior. If these results are compared to PSE in immobile conditions, whether the approach behavior makes it easier to recognize stimuli as happy or whether the avoidance behavior makes it easier to recognize stimuli as fearful should be clarified.

Third, neutral expressions help assess which facial expression is affected by a given behavior. Each experiment compared two expressions. Experiment 1 could not distinguish whether anger recognition was promoted or whether smile recognition was suppressed when the PSE decreased. In addition, the behavior that associates facial expressions can change with approach or avoidance, depending on another facial expression being compared [4]. For example, anger is quicker to avoid compared to happiness, but quicker to approach when compared to fear. Comparing a neutral expression with another expression can help verify whether a given behavior exerted an effect solely based on that expression.

Fourth, we did not verify whether participants identified the stimuli as the intended facial expressions, such as anger, fear, and happiness. Even in the case of facial stimuli that include other expressive components and are not recognized as “anger” in an absolute sense, the presence of significant relative changes between approach and avoidance conditions suggests that these actions modify the recognition of “anger.” To confirm if the stimuli were correctly recognized, it would be beneficial to test when participants freely respond to the model’s expressions to ascertain if they are perceived as the intended expressions. Thus, we could observe an effect by approach–avoidance behavior not only in the one dimension between the two facial expressions used in the experiment but also from a multi-dimensional approach by including other facial expressions.

Finally, the facial models used in this study might have been more likely to appear positive overall. One factor that contributed to facial expression being recognized as positive in Experiment 1 was the greater distance. However, the possibility that the face models themselves had positive facial expressions cannot be denied. Therefore, it is desirable to verify result reproducibility by regenerating a new model.

Future research should clarify the mechanism driving the relationship between approach– avoidance behavior and facial expression recognition. When performing approach–avoidance behaviors, visual information and bodily sensations simultaneously change because of physical movement. Our results are attributable to either visual information or bodily sensations affecting facial expression recognition. To verify which cue is dominant for this effect, a video that enhances vection can be presented, wherein the participant perceives approaching or avoiding while maintaining their somatic sensation. If this condition presents the same results as those in this study, visual information can be confirmed as an essential cue driving cognitive processing for facial expression recognition. Alternatively, bodily approach–avoidance movements can be synchronized with a model in the VR space, such that the model moves as much as participants to retain their visual information.

Moreover, other forms of cognition, such as intimacy [31] and trust [39,40], affect approach– avoidance behaviors. If a reverse relationship is found, as in this study, these other forms of cognition would also be affected by approach–avoidance behaviors. Further, all participants were men; therefore, future research should recruit women.

## Conclusion

We conducted three psychophysical experiments combining approach–avoidance behavior and facial expression recognition to investigate how behavior affects facial expression recognition. Our results indicate that a face is recognized as positive when approaching and as negative when avoiding. Additionally, this relationship is changed to a face that is recognized as an angry, threatening facial expression when approaching for a close distance. These findings suggest that approach–avoidance behaviors are strongly connected to facial expression recognition owing to unconscious learning rooted in biological instincts.

## Acknowledgments

This work was supported by JSPS KAKENHI (Grant Numbers JP21K21315 and JP22K17987 to H.T., JP20H05956 to S.N., and JP20H04273 to T.M.).

## Author contributions

All authors conceived the experiments. Y.K. conducted the experiments. H.T., S.N., and T.M. supervised the study. Y.K. and H.T. interpreted the results and wrote the manuscript’s initial draft. All authors approved the final version of the manuscript.

## Data availability

The datasets generated and/or analyzed during the current study are available from the corresponding author upon reasonable request.

